# On the Origin of Ultraslow Spontaneous Na^+^ Fluctuations in Neurons of the Neonatal Forebrain

**DOI:** 10.1101/2020.05.29.123026

**Authors:** Carlos Perez, Lisa Felix, Christine R. Rose, Ghanim Ullah

**Affiliations:** Department of Physics, University of South Florida, Tampa, FL 33620, USA; Institute of Neurobiology, Faculty of Mathematics and Natural Sciences, Heinrich Heine University Düsseldorf, 40225 Düsseldorf, Germany

## Abstract

Spontaneous neuronal and astrocytic activity in the neonate forebrain is believed to drive the maturation of individual cells and their integration into complex brain-region-specific networks. The previously reported forms include bursts of electrical activity and oscillations in intracellular Ca^2+^ concentration. Here, we use ratiometric Na^+^ imaging to demonstrate spontaneous fluctuations in the intracellular Na^+^ concentration of CA1 pyramidal neurons and astrocytes in tissue slices obtained from the hippocampus of mice at postnatal days 2-4 (P2-4). These occur at very low frequency (∼2/h), can last minutes with amplitudes up to several mM, and mostly disappear after the first postnatal week. To further study the mechanisms that may generate such spontaneous fluctuations in neurons, we model a network consisting of pyramidal neurons and interneurons. Experimentally observed Na^+^ fluctuations are mimicked when GABAergic inhibition in the simulated network is inverted. Both our experiments and computational model show that the application of tetrodotoxin to block voltage-gated Na^+^ channels or of inhibitors targeting GABAergic signaling respectively, significantly diminish the neuronal Na^+^ fluctuations. On the other hand, blocking a variety of other ion channels, receptors, or transporters including glutamatergic pathways, does not have significant effects. In addition, our model shows that the amplitude and duration of Na^+^ fluctuations decrease as we increase the strength of glial K^+^ uptake. Furthermore, neurons with smaller somatic volumes exhibit fluctuations with higher frequency and amplitude. As opposed to this, the larger relative size of the extracellular with respect to intracellular space observed in neonatal brain exerts a dampening effect. Finally, our model also predicts that these periods of spontaneous Na^+^ influx leave neonatal neuronal networks more vulnerable to hyperactivity when compared to mature brain. Taken together, our model thus confirms the experimental observations, and offers additional insight into how the neonatal environment shapes early signaling in the brain.

**Author Summary:** Spontaneous neuronal and astrocytic activity during the early postnatal period is crucial to the development and physiology of the neonate forebrain. Elucidating the origin of this activity is key to our understanding of the cell maturation and formation of brain-region-specific networks. This study reports spontaneous, ultraslow, large-amplitude, long-lasting fluctuations in the intracellular Na^+^ concentration of neurons and astrocytes in the hippocampus of mice at postnatal days 2-4 that mostly disappear after the first postnatal week. We combine ratiometric Na^+^ imaging and pharmacological manipulations with a detailed computational model of neuronal networks in the neonatal and adult brain to provide key insights into the origin of these Na^+^ fluctuations. Furthermore, our model predicts that these periods of spontaneous Na^+^ influx leave neonatal neuronal networks more vulnerable to hyperactivity when compared to mature brain.

## Introduction

Spontaneous neuronal activity is a hallmark of the developing central nervous system [1], and has been described in terms of intracellular Ca^2+^ oscillations both in neurons and astrocytes [2-5] and bursts of neuronal action potentials [6-8]. This activity is believed to promote the maturation of individual cells and their integration into complex brain-region-specific networks [1, 9-11]. In the rodent hippocampus, early network activity and Ca^2+^ oscillations are mainly attributed to the excitatory role of GABAergic transmission originating from inhibitory neurons [7, 12-14].

The excitatory action of GABAergic neurotransmission is one of the most notable characteristics that distinguish neonate brain from the mature brain, where GABA typically inhibits neuronal networks [1, 7, 8, 10-12, 15-17]. While recent work has also called the inhibitory action of GABA on cortical networks into question [18], there are many other pathways that could play a significant role in the observed spontaneous activity in neonate brain (discussed below). Additional key features of the early network oscillations in the hippocampus include their synchronous behavior across most of the neuronal network, modulation by glutamate, recurrence with regular frequency, and a limitation to early post-natal development [2, 7, 12].

More recently, Felix and co-workers [5] reported a new form of seemingly spontaneous activity in acutely isolated tissue slices of hippocampus and cortex of neonatal mice. It consists of spontaneous fluctuations in intracellular Na^+^ both in astrocytes and neurons, which occur in ∼25% of pyramidal neurons and ∼40% of astrocytes tested. Na^+^ fluctuations are ultraslow in nature, averaging ∼2 fluctuations/hour, are not synchronized between cells, and are not significantly affected by an array of pharmacological blockers for various channels, receptors, and transporters. Only using the voltage-gated Na^+^ channel (VGSC) blocker tetrodotoxin (TTX) diminished the Na^+^ fluctuations in neurons and astrocytes, indicating that they are driven by the generation of neuronal action potentials. In addition, neuronal fluctuations were significantly reduced by the application of the GABA_A_ receptor antagonist bicuculline, suggesting the involvement of GABAergic neurotransmission.

This paper follows up on the latter study [5], and uses dual experiment-theory approach to systematically confirm, and further investigate the properties of neuronal Na^+^ fluctuations in the neonate hippocampal CA1 area and to identify the pathways that generate and shape them. Notably, a range of factors that play a key role in controlling the dynamics of extra- and intracellular ion concentrations, are not fully developed in the neonate forebrain [13, 19-22]. These factors, such as the cellular uptake capacity of K^+^ from the extracellular space (ECS), the expression levels of the three isoforms (α1, α2, and α3) of the Na^+^/K^+^ pump that restore resting Na^+^ and K^+^ concentrations, the ratio of intra- to extracellular volumes, and the magnitude of relative shrinkage of the ECS in response to neuronal stimulus, all increase with age and cannot be easily manipulated experimentally [19]. The gap-junctional network between astrocytes is also less developed in neonates and therefore has a lower capacity for the spatial buffering of ions, neurotransmitters released by neurons, and metabolites [19, 21]. At the same time, the synaptic density and expression levels of most isoforms of AMPA and NMDA receptors are very low in neonates and only begin to increase rapidly during the second week [13]. Additionally, while GABAergic synapses develop earlier than their glutamatergic counterparts, synaptogenesis is incomplete and ongoing. Therefore, synapses of varying strengths exist across the network. Each of these aspects impacts the others and their individual specific roles in the early spontaneous activity is consequently difficult to test experimentally. Their involvement in neonatal Na^+^ fluctuations will therefore be addressed for the first time by the data-driven modelling approach here.

We employ ratiometric Na^+^ imaging in tissue slices of the hippocampal CA1 region obtained from neonate animals at postnatal days 2-4 (P2-4) and juveniles at P14-21 to record intracellular Na^+^ fluctuations in both age groups. We begin by reporting the key statistics about spontaneous Na^+^ fluctuations observed in neonates and juveniles. Next, we develop a detailed network model, consisting of pyramidal cells and inhibitory neurons, which also incorporates the exchange of K^+^ in the ECS with astrocytes and perfusion solution *in vitro* (or vasculature in intact brain). Individual neurons are modeled by Hodgkin-Huxley type formalism for membrane potential and rate equations for intra- and extracellular ion concentrations. In addition to closely reproducing our experimental results, the model provides new key insights into the origin of spontaneous slow Na^+^ oscillations in neonates. Furthermore, our model also predicts that the network representing a developing brain is more hyperexcitable when compared to mature brain.

## Results

### Pyramidal neurons in neonate hippocampus exhibit spontaneous ultraslow Na^+^ fluctuations

Acutely isolated parasagittal slices from hippocampi of neonatal mice (P2-4) were bolus-stained with the sodium-sensitive ratiometric dye SBFI-AM along the CA1 region (Figure 1A1). Experimental measurements lasted for 60 minutes, with an imaging frequency of 0.2 Hz. Astrocytes were identified via SR101 staining (Figure 1A1), and were analyzed separately to the neurons in the pyramidal layer. Out of the measured cells, 26% of neurons (n=63/243) and 38% of astrocytes (n=36/97) showed detectable fluctuations in their intracellular Na^+^ concentrations (Figure 1A2, 1B). Detection threshold was calculated individually for each cell, and was defined as being 3 times the standard deviation of the baseline noise of each ROI analyzed (this ranged from 0.28 to 2.04 mM). Astrocyte Na^+^ fluctuations were 10.3 ± 0.7 minutes long, at a frequency of 1.3 ± 0.2 signals/hour and with average amplitudes of 2.4 ± 0.2 mM. Neuronal Na^+^ fluctuations had an average duration of 8.6 ± 0.4 minutes. They occurred at a frequency of 2 ± 0.2 fluctuations/hour with average amplitudes of 2.7 ± 0.12 mM. The high variability in the shapes of fluctuations is demonstrated in Figure 1A2. Apparent synchronicity between cells of the same or different classes was only observed rarely, confirming the observations reported in our earlier study [5].

**Figure 1.**
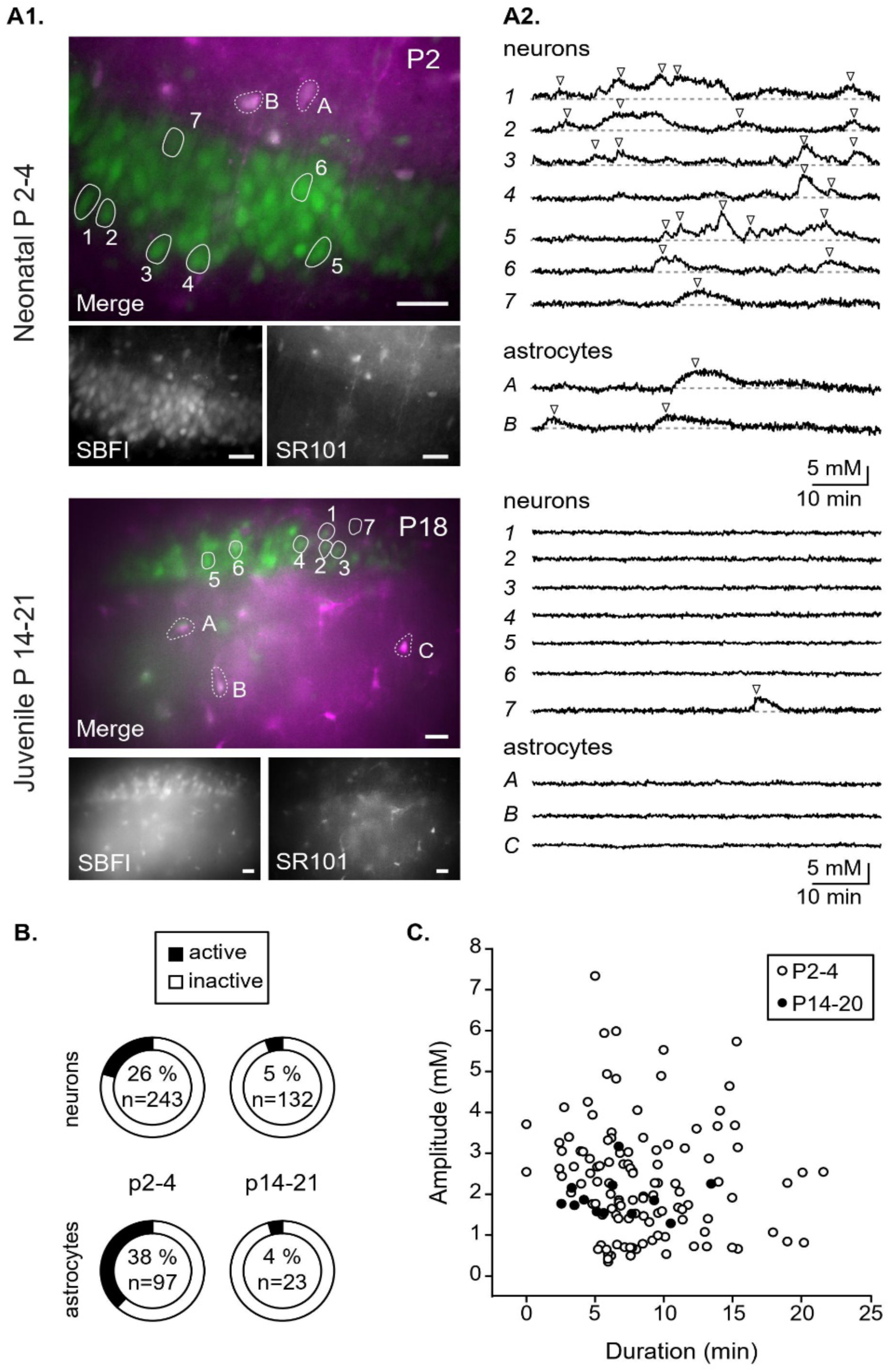
In situ experiments. (A1) Images showing representative stainings in the CA1 region of the neonatal (P4; upper images) and juvenile (P18; lower images) hippocampus. In the merge, SBFI is shown in green and SR101 in magenta. ROIs representing cell bodies of neurons and astrocytes are labeled with numbers and letters, respectively. Scale bars: 20 µm. (A2) Na^+^ fluctuations in the ROIs as depicted in (A1). (B) The percentage of pyramidal neurons and astrocytes showing activity for each age group and the total number of cells measured. (C) Scatter plot showing the peak amplitude and duration of neuronal fluctuations within the two indicated age groups.

To investigate the developmental profile of the fluctuations, the same protocol was repeated in hippocampal tissue from juvenile (P14-20) mice. Here, only 5.3% of all measured neurons (n=7/132) and 4.3% of all measured astrocytes (n=1/23) showed fluctuations in their intracellular Na^+^ concentrations (Figure 1B). This strong reduction confirmed the significant down-regulation of spontaneous Na^+^ oscillations from neonatal to juvenile animals reported recently [5]. However, the properties of the neuronal fluctuations themselves remained unchanged during postnatal development, with the average amplitude, frequency, and duration being 1.9 ± 0.13 mM, 2 ± 0.3 fluctuations/hour, and 6.5 ± 0.9 minutes in juvenile tissue (Figure 1C).

### Spontaneous Na^+^ fluctuations are reproduced by a computational model with excitatory GABAergic neurotransmission

To explore the properties and mechanisms of neonate neuronal Na^+^ fluctuations, we developed a computational model consisting of CA1 pyramidal cells and inhibitory neurons as detailed in the Methods section. Resulting typical time traces of intracellular Na^+^ from four randomly selected excitatory neurons in a network representative of the juvenile hippocampus (where GABAergic neurotransmission is inhibitory) are shown in the right panel of Figure 2A. Na^+^ in individual neurons shows minor irregular fluctuations of less than 0.05 mM around the resting values mostly because of the random synaptic inputs from the network. However, no clear large-amplitude fluctuations can be seen in the network. To mimic neonates, we invert the sign of I-to-E and I-to-I synaptic inputs, making the GABAergic neurotransmission excitatory. The inverted inhibition results in the occurrence of spontaneous Na^+^ fluctuations in the low mM range in individual neurons that persist for several minutes (Figure 2A, left column). In some cases, the peak amplitude of oscillations reached values of more than 5 mM.

**Figure 2:**
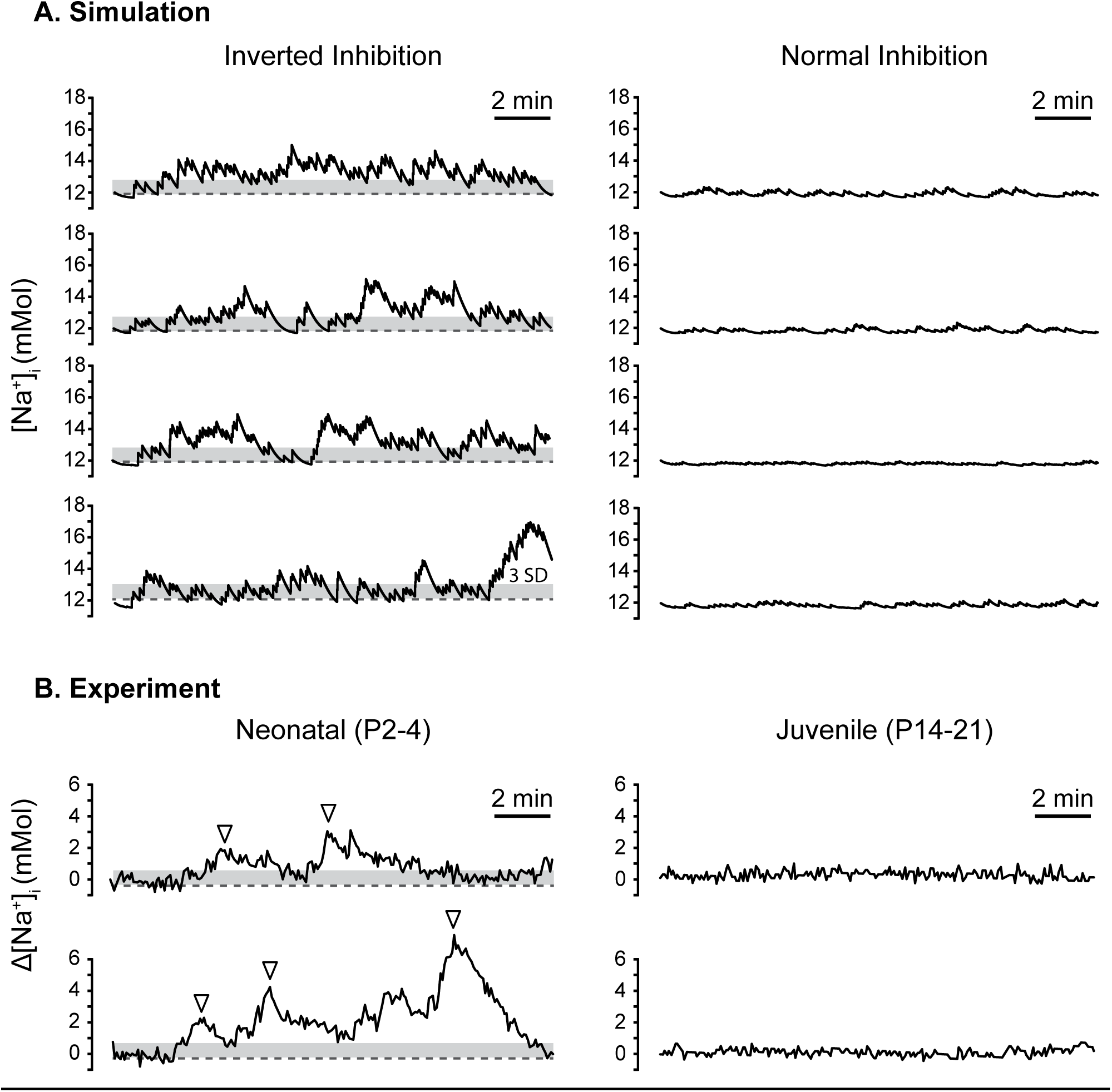
Simulated spontaneous activity in 4 example neurons with excitatory GABAergic neurotransmission representing neonatal hippocampus (A, left) and mature inhibition representing juvenile hippocampus (A, right). Grey bar indicates three times the average standard deviation in experimental traces upwards of the mean. (B) Experimental data, showing excerpts from example measurements shown in Figure 1, both from neonatal neurons (P2-4, left; cell 3- upper; cell 5- lower), and juvenile neurons (P14-21, right; cell 1- upper; cell 2- lower). Traces show changes in intracellular Na^+^ concentration over 17 minutes, a time course directly comparable to (A).

The simulated data shows a comparable pattern of irregular fluctuations to the experimental results (Figure 2B). The properties of these events are very similar—with peak amplitudes mostly in the 2-3 mM range and durations spanning over several minutes. However, the simulated data also appears to show a high rate of low amplitude spiking, apparently absent from the experimental traces. As mentioned above, the detection threshold for experimental data ranged from 0.28 to 2.04 mM (see also Figure 2B), and the imaging frequency was kept at 0.2 Hz in order to prevent phototoxic effects during the long-lasting continuous recordings. Fast, low amplitude transients as revealed in simulated experiments are thus below the experimental detection threshold-as indicated in Figure 2.

### Neonate network does not exhibit spontaneous fluctuations in [K^+^]_o_

Since the dynamics of Na^+^ and K^+^ are generally coupled in mature brain, we next look at K^+^ concentration in the ECS of individual neurons ([K^+^]_o_) in the network to see if it exhibits similar spontaneous fluctuations. A sample trace for a randomly selected neuron is shown in Figure 3A (gray). As clear from the figure, there are only minimal fluctuations in [K^+^]_o_ (peak amplitudes of residual changes are < 0.05 mM) with respect to the resting state when compared to the much larger [Na^+^]_i_ fluctuations in the same cell (gray line in Figure 3B). Next, we recorded [K^+^]_o_ traces for all pyramidal neurons in the network and calculated the mean [K^+^]_o_ (averaged over all excitatory neurons). The mean [K^+^]_o_ as a function of time shows that all excitatory neurons in the network exhibit very small changes in the [K^+^]_o_, which are essentially canceled out at the network level (Figure 3A, black line). The mean intracellular Na^+^ fluctuates slightly more than the mean [K^+^]_o_ (Figure 3B, black line). However, a comparison between the traces showing the average Na^+^ over all excitatory neurons in the network and that from the single neuron indicates that the amplitude of Na^+^ fluctuations varies from cell to cell and that they are not necessarily phase-locked. All these observations are in agreement with experimental results reported above.

**Figure 3:**
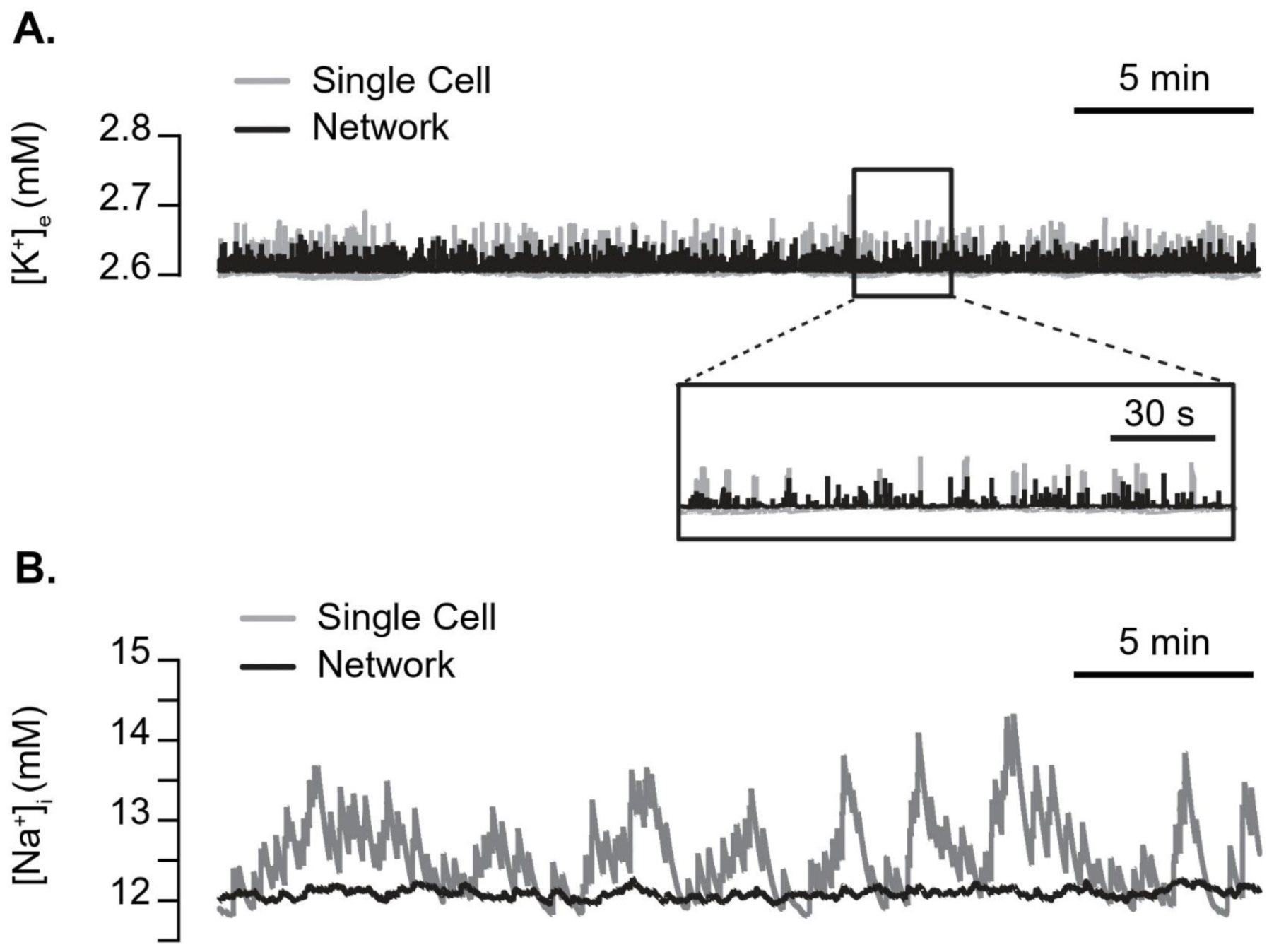
Simulated spontaneous fluctuations in intracellular Na^+^ ([Na^+^]_i_) are not coupled with significant fluctuations in extracellular K^+^ ([K^+^]_o_). [K^+^]_o_ (A) and [Na^+^]_i_ (B) time traces from a randomly selected excitatory neuron (gray) and averaged over the entire excitatory network (black).

### The model replicates the observed effects of TTX and other blockers

We next performed imaging experiments in which various blockers were applied. Addition of 0.5 µM TTX reduced the number of neurons showing fluctuations to 4 % (n=7/167), suggesting a dependence on action potential generation via the opening of voltage-gated Na^+^ channels (Figure 4A). However, blocking of glutamatergic receptors with a cocktail containing APV (100 µM), NBQX (25 µM), and MPEP (25 µM) (targeting NMDA, AMPA/kainate, and mGluR5 receptors, respectively) had no effect on the number of neurons showing fluctuations (21% active, n=33/155) (Figure 4A). Additionally, the role of GABAergic signaling was tested via combined application of bicuculline (10 µM), CGP-55845 (5 µM), NNC-711 (100 µM), and SNAP-5114 (100 µM) (antagonists for GABA_A_ receptors, GABA_B_ receptors, GABA transporters GAT1, and GAT2/3, respectively). This combination of antagonists reduced the number of active neurons to a similar degree as TTX (3% active, n=5/158) (Figure 4A). These data are concordant with the results previously published [5], and suggest that the slow fluctuations in intracellular Na^+^ are produced by the accumulation of Na^+^ during trains of action potentials, triggered by GABAergic transmission.

**Figure 4:**
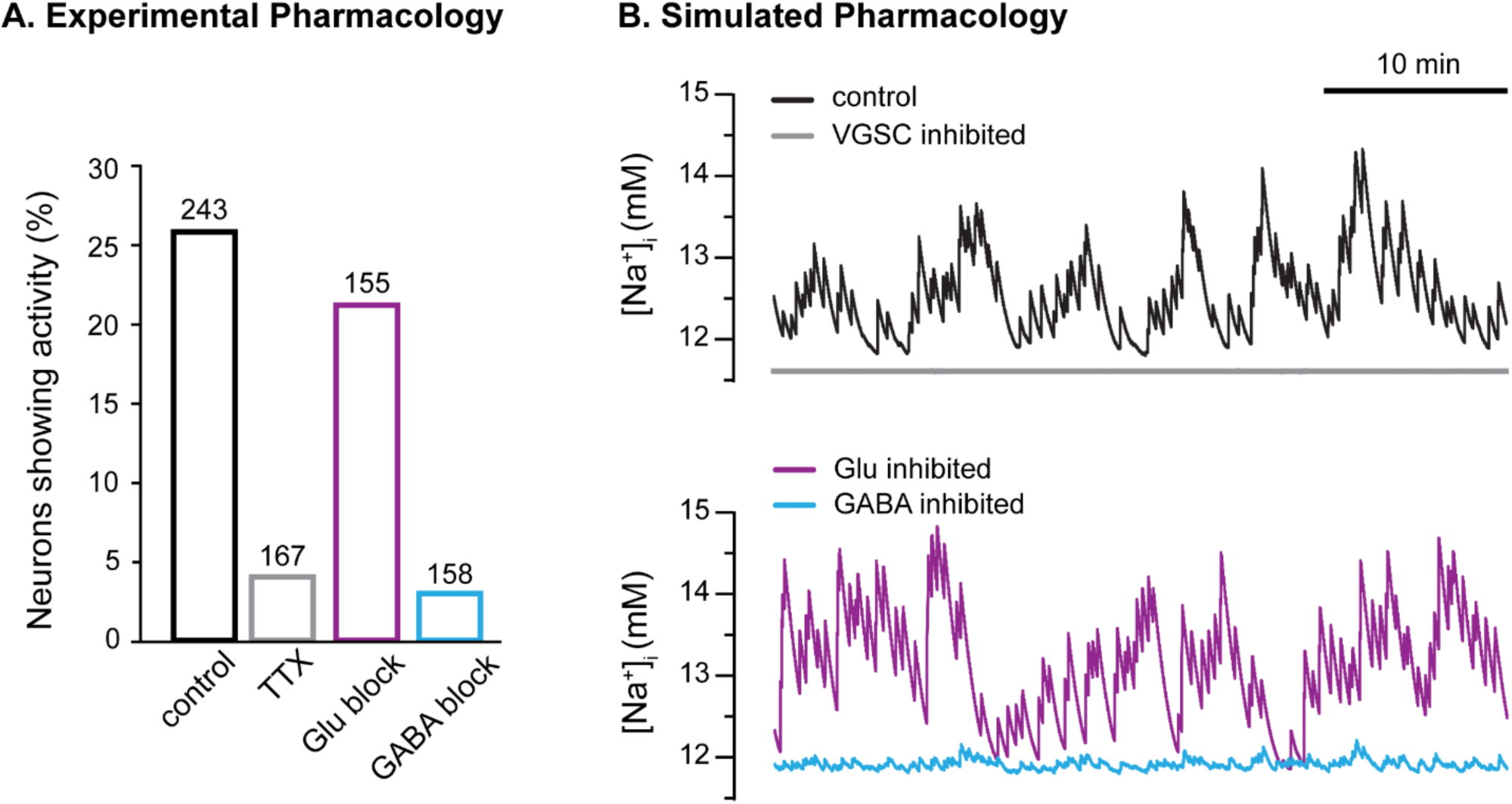
Inhibiting GABA_A_ receptors or voltage-gated Na^+^ channels eliminates [Na^+^]_i_ fluctuations, whereas blocking glutamatergic synaptic inputs has little effect. (A) Bar plot showing the percentage of neurons exhibiting Na^+^ fluctuations as determined in experiments under the four conditions simulated in (B). That is, the percentage of neurons exhibiting Na^+^ fluctuations in slices from juveniles under control conditions (black) and in the presence of 0.5 µM TTX to block voltage gated Na^+^ channels (gray), a cocktail containing APV (100 µM), NBQX (25 µM), and MPEP (25 µM) to block glutamatergic receptors (purple), and a combined application of bicuculline (10 µM), CGP-55845 (5 µM), NNC-711 (100 µM), and SNAP-5114 (100 µM) to block GABAergic signaling (cyan). (B) Time trace of [Na^+^]_i_ from a randomly selected excitatory neuron in the network in control conditions (inverted inhibition, representing neonatal brain) (black, top panel), with voltage-gated Na^+^ channels blocked (gray, top panel), glutamatergic synapses blocked (purple, bottom panel), and GABAergic synapses blocked (cyan, bottom panel).

The pharmacological profile of the experimentally observed Na^+^ fluctuations in the neonatal brain summarized above strongly suggests that the excitatory effect of GABAergic neurotransmission plays a key role in their generation, whereas glutamatergic activity contributes very little. Before making model-based predictions, we first confirm that our model reproduces these key observations in our experiments. We first incorporate the effect of TTX in the model by setting the peak conductance of voltage-gated Na^+^ channels to zero. We also mimic the effect of blocking ionotropic glutamate receptors with CNQX and APV by setting E-to-E and E-to-I synaptic conductances to zero. Finally, we mimic the effect of blocking GABAergic transmission on the activity of the network, and set the I-to-I and I-to-E synaptic currents to zero, thereby removing all GABA_A_-receptor-related effects. The model results are largely in line with observations, where we see that inhibiting GABA-related currents and voltage-gated Na^+^ channels mostly eliminate Na^+^ fluctuations and blocking NMDA and AMPA synaptic inputs has little effect on the observed spontaneous activity (Figure 4B).

### Spontaneous Na^+^ fluctuations are shaped by neuronal morphology and glial K^+^ uptake capacity

As pointed out above, significant changes occur in the physical and functional properties of the neurons during postnatal maturation at the synaptic, single cell, and network levels [13, 19]. Therefore, we use the model to examine if changes in some key physical and functional characteristics of the network such as the neuronal radius (r_in_), the ratio of ICS to ECS (β), and glial K^+^ uptake rate play any role in the observed Na^+^ fluctuations. In the following, we show Na^+^ time traces for four randomly selected excitatory neurons. We observe that smaller neurons in general exhibit larger Na^+^ fluctuations (p>0.001, Figure 5A, left panels). Both the amplitude and frequency of fluctuations decrease as we increase r_in_ (Figure 5A, center panels). The panel on the right in Figure 5A (and Figure 5B, C) shows the average amplitude of Na^+^ fluctuations as we change the parameter of interest.

**Figure 5:**
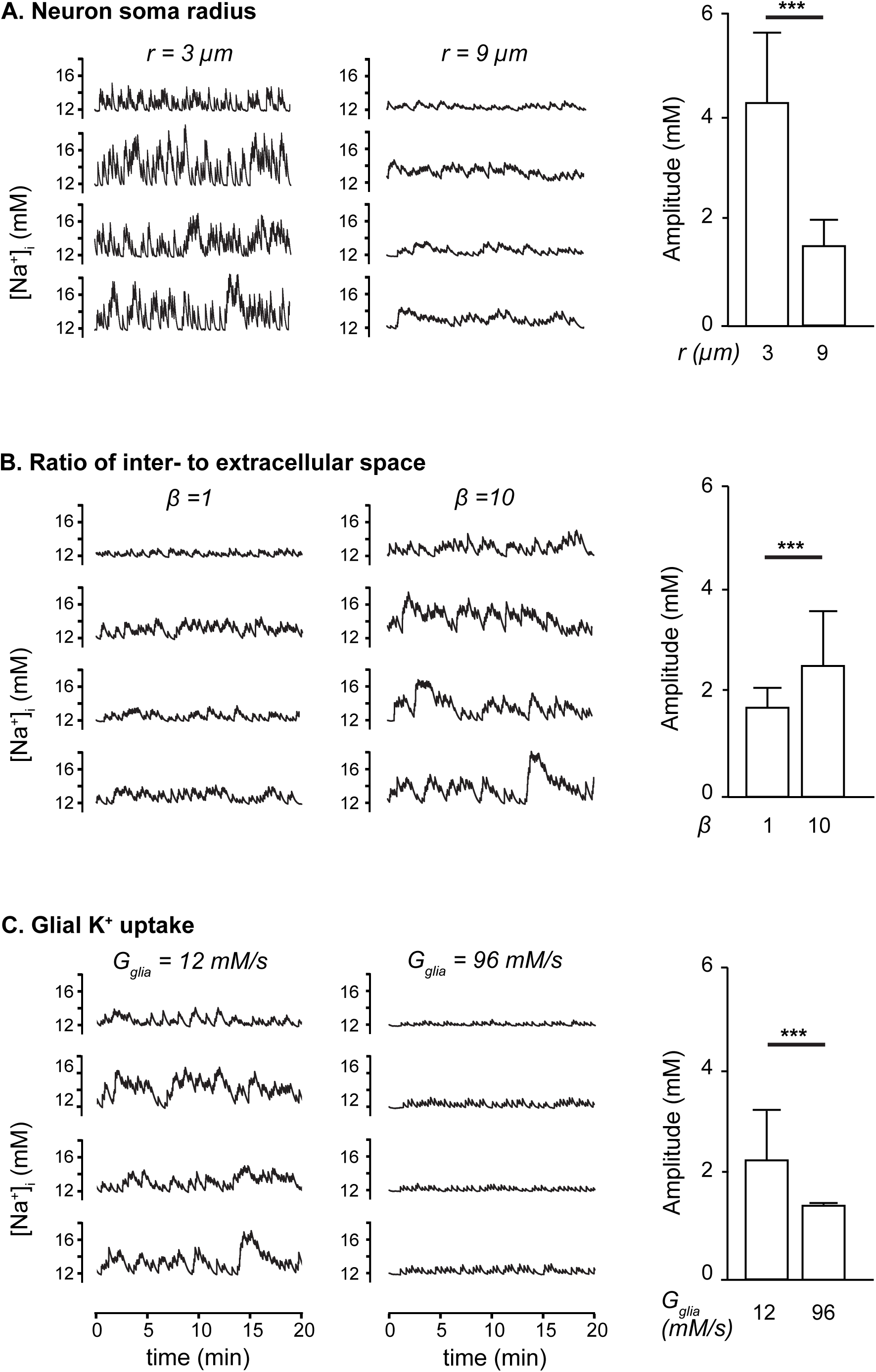
The neuronal radius, ratio of ECS to ICS (β), and K^+^ uptake capacity of glia affect spontaneous Na^+^ fluctuations. (A) Time traces of [Na^+^]_i_ for five excitatory neurons from a network representing neonatal brain with a neuronal radius of 3 μm (left panels) and 9 μm (center panels). The panel on the right shows the mean amplitude of Na^+^ fluctuations (averaged over all pyramidal neurons in the network) under the two conditions. The error bars indicate the standard deviation of the mean. β was fixed at 2.5. (B) Same as (A) at β = 1 (left panels) and 10 (center panels). (C) Same as (A) with maximum glial K^+^ buffering strength G_glia_=12 mM/s (left panels) and G_glia_=96 mM/s (right panels). The radius of individual neurons is set at 6 μm in both (B) and (C). ***: p>0.001.

The observed fraction of ECS with respect to ICS in neonates is approximately 40% (β = 2.5) [23, 24], compared to adult animals where ECS is about 15% of the ICS (β ∼ 7) [25, 26]. We vary β from 1 to 10 to see how it affects Na^+^ fluctuations. An opposite trend as compared to neuronal radius can be seen when we change β, where larger β results in Na^+^ fluctuations that are larger in amplitude and have longer duration (Figure 5B, center panels) compared to those in neurons with smaller β values (p>0.001, Figure 5B, left panels). Thus the relative larger ECS in neonates does not favor the generation of large Na^+^ fluctuations, but on the contrary dampens ion changes.

The expression levels of astrocytic channels and transporters involved in K^+^ uptake (Na^+^/K^+^ ATPase, Kir4.1 channels, and Na^+^/K^+^/Cl^-^ co-transporter 1 (NKCC1)) and connexins forming gap junctions are low in neonates [20, 21]. Astrocytes in the neonate brain, therefore, have a lower capacity for uptake of extracellular K^+^ released by neurons [19]. To analyze the influence of glial K^+^ uptake, we varied the maximum glial K^+^ uptake strength in the model from 12 mM/sec (significantly lower than 66 mM/sec - the value used for mature neurons in [27]) to 96 mM/sec to see how it affects Na^+^ fluctuations. We observed a strong effect of varying peak glial K^+^ uptake on the amplitude and frequency of Na^+^ fluctuations (p>0.001). Overall, the amplitude and frequency of Na^+^ fluctuations decrease as we increase peak glial K^+^ uptake (Figure 5C).

### The model predicts a higher propensity of neonatal brain for hyperexcitability

Significant evidence shows that the neonatal brain is more hyperexcitable [28-31]. For example, the frequency of seizure incidences is highest in the immature human brain [15, 32, 33]. Critical periods where the animal brain is prone to seizures have also been well-documented [15]. Various epileptogenic agents and conditions, including an increase in [K^+^]_o_, result in sigmoid-shaped age-dependence of seizure susceptibility in postnatal hippocampus [15, 34-36]. The developmental changes in GABAergic function are suspected to play a key role in the change in seizure threshold and the higher incidences of seizures in neonates [16, 37].

To test this hypothesis, we next investigate how excitatory GABAergic neurotransmission affects the excitability of the network in response to different levels of [K^+^]_o_. In the model, we take the average frequency of action potential (AP) generation (average number of spikes per minute per neuron) of all excitatory neurons as a measure of the susceptibility of the network to hyperexcited states such as seizures. As illustrated in Figure 6A, the overall AP frequency is significantly larger in the network with inverted inhibition (representing the neonatal brain) than the network with mature inhibition (representing a mature network). For all [K^+^]_o_ values tested, the average AP frequency in the neonatal network is doubled that of mature network. Thus, our simulation predicts that inverted inhibition strongly increases the excitability of neurons, indicating a significantly lower threshold for hyperexcitability in neonates (Figure 6). Our simulations also show that decreasing the radius of neurons or the K^+^ uptake capacity of astrocytes further increases the vulnerability of neonate brain to hyperactivity (not shown).

**Figure 6:**
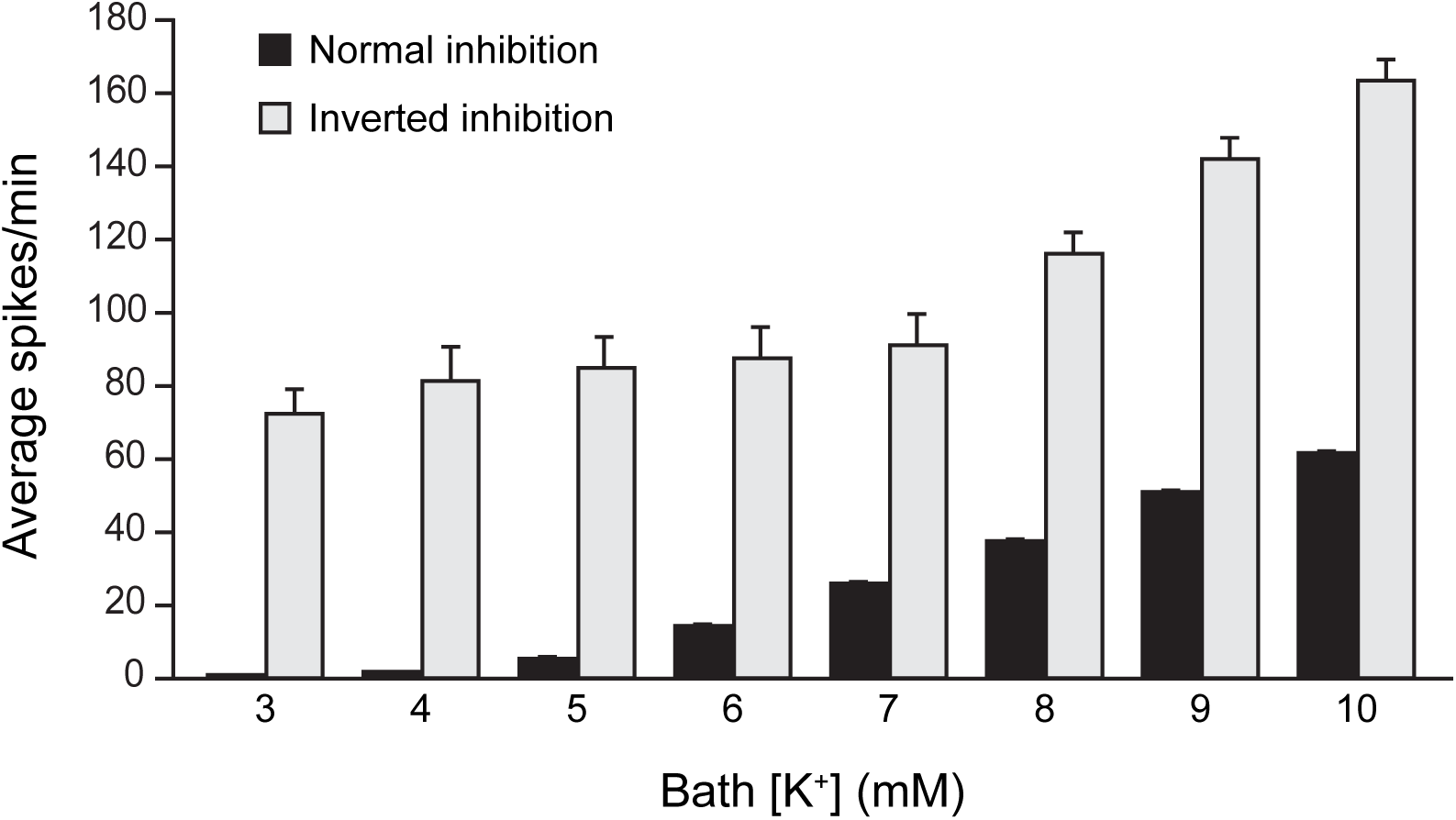
Inverted inhibition leaves the network more prone to hyperactivity. (A) Bar plot showing the number of spikes per minute averaged over all excitatory neurons as we systematically increase K^+^ concentration in the bath. The black and gray bars correspond to neural network with mature and inverted inhibition respectively. The error bars indicate the standard deviation of the mean.

## Discussion

Spontaneous neuronal and astrocytic activity is the hallmark of the developing brain and drives cell differentiation, maturation, and network formation [1-9] [10, 11]. In the neonate hippocampus, this activity is mostly attributed to the excitatory effect of GABAergic neurotransmission [14]. While spontaneous activity has also been shown in cortical neuronal networks, these appear to originate primarily from pace-maker cells in the piriform cortex, and are driven by a separate mechanism involving both glutamate and GABA [38]. In contrast, hippocampal early network oscillations stem solely from GABA released by interneurons. Hippocampal interneurons constitute a diverse group of cells, including the fast-spiking inhibitory neurons simulated in this study. These cells have previously been implicated in the generation of early network activity in the hippocampus and cortex as the timing of their synapse formation around pyramidal cells closely match that of the appearance of giant depolarizing potentials in the neonatal brain. Additionally, the optogenetic blocking of their activity was shown to halt spontaneous giant depolarizing potentials almost entirely [39].

The excitatory effect of GABA on neurons is related to the higher expression of the Na^+^/K^+^/Cl^-^ cotransporter as compared to the K^+^/Cl^-^ cotransporter in the first week after birth. This results in elevated intracellular Cl^-^, leading to an outwardly directed Cl^-^ gradient [40-42], and in an efflux of Cl^-^ when GABA_A_ receptor channels open, causing the post-synaptic neuron to depolarize [18].

In this study, we report spontaneous, ultraslow fluctuations in the intracellular Na^+^ concentration of CA1 pyramidal neurons and astrocytes in tissue slices from mouse hippocampus, recorded using ratiometric Na^+^ imaging, thereby confirming our recent observations [5]. As reported in the latter study, these spontaneous fluctuations are primarily present during the first postnatal week and rapidly diminish afterwards. Unlike the giant depolarizing potentials (GDPs) and early network Ca^2+^ oscillations observed in the hippocampus previously [2, 7, 12], the Na^+^ fluctuations reported here are not synchronous, involve only about a quarter of all pyramidal cells recorded, are not significantly modulated by glutamatergic neurotransmission, and do not occur with regular frequency. Furthermore, these fluctuations are extremely rare (∼2/hour), long-lasting (each fluctuation lasting up to several minutes), and strongly attenuated by the application of TTX to block VGSCs and application of inhibitors of GABAergic neurotransmission. A range of other pharmacological blockers targeting various channels, receptors, co-transporters, or transporters did not significantly affect these fluctuations (Figure 4 and [5]).

To investigate the origin of the spontaneous neuronal Na^+^ fluctuations further, we developed a detailed computational model that represents a hippocampal network, incorporating the three main cell types (pyramidal cells, inhibitory neurons, and astrocytes) and ion concentration dynamics in principal neurons and the extracellular space. In agreement with observations from our experimental data presented here and the earlier experimental study [5], the computational results suggest that voltage-gated Na^+^ channels and the excitatory effect of GABAergic neurotransmission play key roles in the generation of the ultraslow Na^+^ fluctuations. Our simulation results also reveal that these fluctuations occur at the individual neuronal level, are not phase-locked, and are not strictly a network phenomenon, thereby confirming experimental results. Moreover, the fluctuations are confined to intracellular Na^+^ and are not observed in extracellular K^+^, further supporting the conclusion that these fluctuations are a local phenomenon.

Because synaptogenesis is ongoing during the first postnatal week, synapses across the neuronal network display varying strengths. This means that while activity such as GDPs can happen synchronously across populations, individual synapses will experience different levels of Na^+^ influx in response to action potentials. A neuron with a large number of strong synapses from an interneuron would therefore have a larger influx of Na^+^ (considering the depolarizing inhibition in the neonate brain) than neurons with fewer, weaker connections. The pattern of connectivity and variations in GABA release between several interneurons could therefore explain the unusually long, irregular, asynchronous fluctuations seen in individual neurons here, as they might arise from the summation of inputs.

In addition to the outwardly directed Cl^-^ gradient and the excitatory action of GABA, the neonate forebrain in the first week after birth is in a constant state of flux where many functional and morphological changes occur along with the differentiation and maturation of cells and the cellular network [7, 12, 13, 19, 43]. Two of the most significant changes include the still ongoing gliogenesis and astrocyte maturation [44-46]. Immature astrocytes have a reduced glial uptake capacity for K^+^ as well as for glutamate compared to the mature brain [19, 21, 47]. Furthermore, the neonate brain exhibits an increased volume fraction of the ECS [23, 24, 48]. These factors along with the morphological properties of cells, play key roles in ion concentration dynamics. Indeed, we found the behavior of intracellular Na^+^ fluctuations to be strongly reliant on neuronal radius. However, the larger extra- to intracellular volume ratio appears to suppress Na^+^ fluctuations, suggesting that the larger relative ECS observed in neonates does not play a significant mechanistic role in the generation of spontaneous activity. Our model also suggests that increasing glial K^+^ uptake capacity results in decreasing the amplitude and frequency of Na^+^ fluctuations in the individual neurons and thus may play a role in their suppression at later stages of postnatal development.

Convincing evidence shows that the developing brain is more hyperexcitable. This is supported by the significantly higher frequency of seizures in the neonatal brain [15, 32, 33]. The higher occurrence of seizures is primarily attributed to the excitatory effect of GABA [49]. Based on the above analysis, we believe that the inability of astrocytes to effectively take up extracellular K^+^ and morphological changes together with the inverted Cl^-^ gradients leave the developing brain more susceptible to hyperexcitability and epileptic seizures. As a proof of concept, we exposed our model network to increasing concentrations of K^+^ in the bath solution, similar to experimental protocols used to generate epileptiform activity in brain slices. Indeed, we found that the network representing the neonate brain is unable to cope with the elevated extracellular K^+^ concentration efficiently and exhibits hyperactivity as we increase bath K^+^. Decreasing the radius of neurons or the K^+^ uptake capacity of astrocytes further increases the vulnerability of neonate brain to hyperactive behavior (not shown).

To summarize, our dual experiment-theory approach asserts that the ultraslow, long-lasting, spontaneous intracellular Na^+^ fluctuations observed in neonate brain are non-synchronous, not coupled with fluctuations in extracellular K^+^, and only occur in a fraction of neurons (and astrocytes, see Figure 1 and [5]). These fluctuations are most likely due to a combination of factors with the excitatory GABAergic neurotransmission and action potential generation playing dominant roles. In addition, other conditions in the neonate brain such as decreased K^+^ uptake capacity of astrocytes and morphological properties of neurons also play key roles. Furthermore, glutamatergic and other pathways do not seem to make notable contributions to the Na^+^ fluctuations. The combination of factors described above also provides an environment in the neonate brain that is conducive to hyperexcitability and seizure-like states. Thus, the experimental and computational work presented here provides deep insights into this newly observed phenomena and its possible link with hyperexcitability-related pathology in the developing brain.

## Materials and Methods

### Experimental Methods

#### Relevant abbreviations and source of chemicals

**MPEP** (2-Methyl-6-(phenylethynyl)pyridine) from Tocris

**APV** ((2*R*)-amino-5-phosphonovaleric acid; (2*R*)-amino-5-phosphonopentanoate) from Cayman Chemical

**NBQX** (2,3-Dioxo-6-nitro-1,2,3,4-tetrahydrobenzo[f]quinoxaline-7-sulfonamide) from Tocris **CGP-55845** ((2S)-3-[[(1S)-1-(3,4-Dichlorophenyl)ethyl]amino-2-hydroxypropyl](phenylmethyl)phosphinic acid hydrochloride) from Sigma-Aldrich

**NNC-711** (1,2,5,6-Tetrahydro-1-[2-[[(diphenylmethylene)amino]oxy]ethyl]-3-pyridinecarboxylic acid hydrochloride) from Tocris

**SNAP-5114** (1-[2-[tris(4-methoxyphenyl)methoxy]ethyl]-(S)-3-piperidinecarboxylic acid) from Sigma-Aldrich

#### Preparation of tissue slices

This study was carried out in accordance with the institutional guidelines of the Heinrich Heine University Düsseldorf, as well as the European Community Council Directive (2010/63/EU). All experiments were communicated to and approved by the animal welfare office of the animal care and use facility of the Heinrich Heine University Düsseldorf (institutional act number: O52/05). In accordance with the German animal welfare act (Articles 4 and 7), no formal additional approval for the post-mortem removal of brain tissue was necessary. In accordance with the recommendations of the European Commission [50], juvenile mice were first anaesthetized with CO_2_ before the animals were quickly decapitated, while animals younger than P10 received no anesthetics.

Acute brain slices with a thickness of 250 µm were generated from mice (*mus musculus*, Balb/C; both sexes) using methods previously published [51]. An artificial cerebro-spinal fluid (ACSF) containing (in mM): 2 CaCl_2_, 1 MgCl_2_ 125 NaCl, 2.5 KCl, 1.25 NaH_2_PO_4_, 26 NaHCO_3_, and 20 glucose was used throughout all experiments and preparation of animals younger than P10. For animals at P10 or older, a modified ACSF (mACSF) was used during preparation, containing a lower CaCl_2_ concentration (0.5 mM), and a higher MgCl_2_ concentration (6 mM) but being otherwise identical to the normal ACSF. Both solutions were bubbled with 95% O_2_/5% CO_2_ to produce a pH of ∼7.4 throughout experiments, and each had an osmolarity of 308-312 mOsm/l. Immediately after slicing, the slices were transferred to a water bath and incubated at 34°C with 0.5-1 µM sulforhodamine 101 (SR101) for 20 minutes, followed by 10 minutes in 34°C ACSF without SR101. During experiments, slices were continuously perfused with ACSF at room temperature. For experiments utilizing antagonists, these were dissolved in ASCF and bath applied for 15 minutes before the beginning, and subsequently throughout the measurements.

#### Sodium Imaging

Slices were dye-loaded using the bolus injection technique (via use of a picospritzer 3, Parker, Cologne, Germany). The sodium-sensitive ratiometric dye SBFI-AM (sodium-binding benzofuran isophthalate-acetoxymethyl ester; Invitrogen, Schwerte, Germany) was used for detection of Na^+^. SBFI was excited alternatingly at 340 nm (Na^+^-insensitive wavelength) and 380 nm (Na^+^-sensitive wavelength) by a PolychromeV monochromator (Thermo Fisher Scientific, Eindhoven, Netherlands). Emission was collected above 420 nm from defined regions of interest (ROIs) drawn around cell somata using an upright microscope (Nikon Eclipse FN-1, Nikon, Düsseldorf, Germany) equipped with a Fluor 40x/0.8W immersion objective (Nikon), and attached to an ORCA FLASH 4.0 LT camera (Hamamatsu Photonics Deutschland GmbH, Herrsching, Germany). The imaging software used was NIS-elements AR v4.5 (Nikon, Düsseldorf, Germany). For the identification of astrocytes [52], SR101 was excited at 575 nm and its emission collected above 590 nm.

#### Data analysis and statistics

For each ROI, a ratio of the sensitive and insensitive emissions was calculated and analyzed using OriginPro 9.0 software (OriginLab Corporation, Northampton, MA, USA). Changes in fluorescence ratio were converted to mM Na^+^ on the basis of an *in situ* calibration performed as reported previously [53, 54]. A signal was defined as being any change from the baseline, if Na^+^ levels exceeded 3 standard deviations of the baseline noise. Each series of experiments was performed on at least four different animals, with ‘n’ reflecting the total number of individual cells analyzed. Values from experiments mentioned in the text are presented as mean ± standard error, while values taken from models are presented as mean ± standard deviation.

### Computational Methods

The basic equations for the membrane potential of individual neurons, various ion channels, and synaptic currents used in our model are adopted from Ref. [55]. The network topology follows the scheme for hippocampus from the same work. As shown in Figure 7, the network consists of pyramidal cells and fast-spiking interneurons with five to one ratio. The results reported in this paper are from a network with 25 excitatory and 5 inhibitory neurons. Astrocytes are not explicitly illustrated as cellular entities in Figure 7, but included in the model through their ability to take up K^+^. Of note, increasing the network size does not change the conclusions from the model (not shown). Each inhibitory neuron makes synaptic connections with 5 adjacent postsynaptic pyramidal neurons (I-to-E synapses). Thus five excitatory and one inhibitory neurons constitute one “domain”. As shown in the “Results” section, we observed significant variability in the neuronal behavior. Approximately 25% of neurons tested exhibited Na^+^ fluctuations. Furthermore, the amplitude, duration, and frequency of the fluctuations varied over a wide range, pointing towards a heterogeneity in the network topology. To incorporate the observed variability in the neuronal behavior, the synaptic strengths vary randomly from one domain to another. For inhibitory-to-inhibitory (I-to-I), excitatory-to-excitatory (E-to-E), and excitatory-to-inhibitory (E- to-I) synapses, we consider all-to-all connections. However, restricting these synapses spatially does not change the conclusions in the paper. We remark that if one wishes to use a network of a different size with all-to-all connections, the maximum strength of these three types of synaptic inputs will need to be scaled according to the network size.

**Figure 7:**
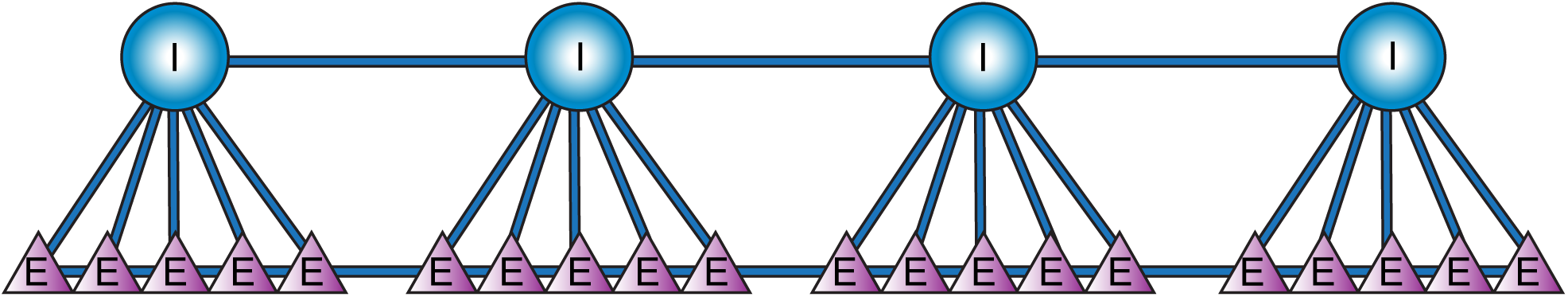
Network schematic showing connections between adjacent neurons within the two neuronal layers. The network consists of pyramidal (E) and inhibitory (I) neurons at five to one ratio, where five excitatory and one inhibitory neurons make one domain. In addition to synaptic inputs, we also consider the diffusion of extracellular K^+^ between neighboring cells. Incorporating Na^+^ and Cl^-^ diffusion in the extracellular space does not change our conclusions (not shown) and is therefore not included in the model.

The equations for individual cells are modified and extended to incorporate the dynamics of various ion species in the intra- and extracellular spaces of the neurons using the formalism previously developed in [27, 56-61]. The change in the membrane potential, *V*_*m*_, for both excitatory and inhibitory neurons in the network is controlled by various Na^+^ (I_Na_), K^+^ (I_K_), and Cl^-^ (I_Cl_) currents, current due to Na^+^/K^+^-ATPase (*I*_*pump*_), and random inputs from neurons that are not a part of the network 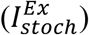, and is given as

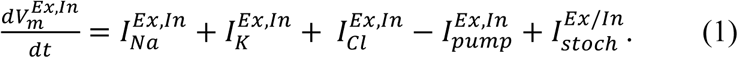

The superscripts *Ex* and *In* correspond to excitatory and inhibitory neurons respectively. The Na^+^ and K^+^ currents consist of active currents corresponding to fast sodium and delayed rectifier potassium channels 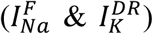, passive leak currents 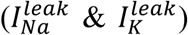, and excitatory synaptic currents 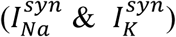. The chloride currents consist of contributions from passive leak current 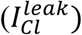 and inhibitory synaptic currents 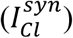.

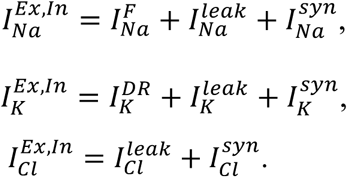

The equations for active neuronal currents are given by the following equations,

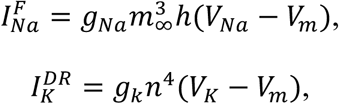

where *g*_*Na*_, *g*_*k*_, *m*_∞_, *h*, and *n* represent the maximum conductance of fast Na^+^ channels, maximum conductance of delayed rectifier K^+^, steady state gating variable for fast Na^+^ activation, fast Na^+^ inactivation variable, and delayed rectifier K^+^ activation variable. As in [55], the gating variables and peak conductances for 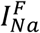, 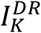, and leak currents for the pyramidal neurons in this study are based on the model of Ermentrout and Kopell [62], which is a reduction of a model due to Traub and Miles [63]. The equations for fast-spiking inhibitory neurons are taken from the model in [64] and [65], which is a reduction of the multi-compartmental model described in Ref. [66]. These equations were originally chosen such that the model would result in the intrinsic frequency as a function of stimulus strength observed in pyramidal cells and fast-spiking inhibitory neurons respectively. The gating variables obey the following equations,

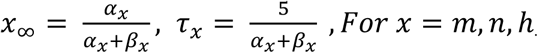

Here x_∞_ and τ_x_ represent the steady state and time constant of the gating variable respectively. The forward and reverse rates (α_x_ and β_x_) for the channel activation and inactivation are calculated using the equations below.

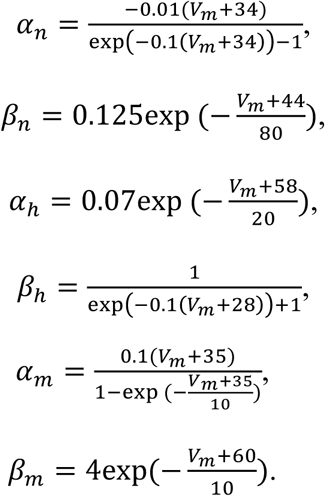

The leak currents are given by

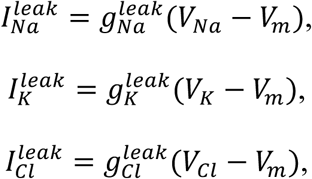

where *V*_*Na*_, *V*_*K*_, and *V*_*Cl*_ are the reversal potentials for Na^+^, K^+^, and Cl^-^ currents respectively and are updated according to the instantaneous values of respective ion concentrations.

The functional form of stochastic current 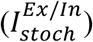 received by each neuron is also based on [55] and is given as

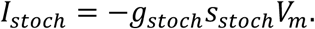

Where *g*_*stoch*_ represents the maximal conductance associated with the stochastic synaptic input and is set to 1 for both cell types. The gating variable s_stoch_ decays exponentially with time constant τ_stoch_= 100 ms during each time step Δ*t*, that is

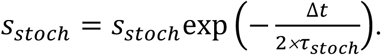

At the end of each time step, s_stoch_ jumps to 1 with probability Δt × *f*_stoch_/1000, where *f*_stoch_ is the mean frequency of the stochastic inputs. These equations simulate the arrival of external synaptic input pulses from the neurons that are not included in the network [55].

The excitatory and inhibitory synaptic currents corresponding to AMPA, NMDA, and GABA receptors are given by the equations below,

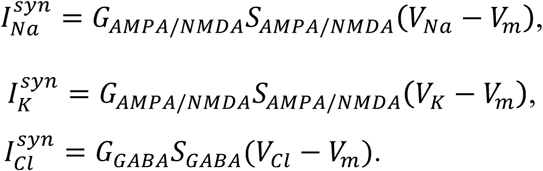

*G*_*AMPA/NMDA*_, *G*_*GABD*_, *S*_*AMPA/NMDA*_, and *S*_*GABD*_ represent the synaptic conductance and gating variables for AMPA and NMDA (represented by a single excitatory current) and GABA receptors. To incorporate the observed variability in neuronal behavior, we randomly select the maximal conductance value for I-to-E synapses inside a single domain from a Gaussian distribution between 0.1 and 3.0 mS/cm^2^. In order to model the excitatory role of GABAergic neurotransmission observed in neonate brain, we change the sign of G_GABA_ from positive to negative.

The change in synaptic gating variables for both excitatory and inhibitory neurons is modeled as in [55]. That is

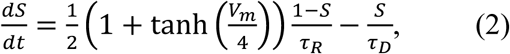

where *τ*_*R*_ and *τ*_*D*_ represent the rise and decay time constants for synaptic signals. The reversal potentials used in the above equations are calculated using the Nernst equilibrium potential equations, i.e.

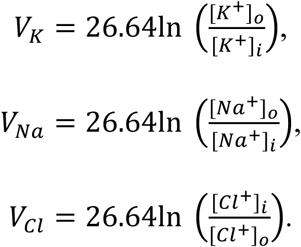

Where [K^+^]_o/i_, [Na^+^]_o/i_, and [Cl^-^]_o/i_ represent the concentration of Na^+^, K^+^, and Cl^-^ outside and inside the neuron respectively. We consider the ECS as a separate compartment surrounding each cell, having a volume of approximately 15% of the intracellular space (ICS) in the hippocampus of adult brain [25, 26] and ∼40% of the ICS in neonates [23, 24]. Each neuron exchanges ions with its ECS compartment through active and passive currents, and the Na^+^/K^+^-ATPase. The ECS compartment can also exchange K^+^ with the glial compartment, perfusion solution (or vasculature in intact brain), and the ECS compartments of the nearby neurons [67-69].

The change in [K^+^]_*o*_ is a function of *I*_*K*_, *I*_*pump*_, uptake by glia surrounding the neuron (*I*_*glia*_), diffusion between the neuron and bath perfusate (*I*_*diff1*_), and lateral diffusion between adjacent neurons (I_diff2_).

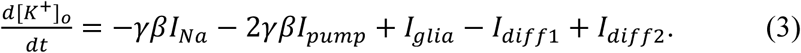

Where *β* is the ratio of ICS to ECS. We set *β* = 7 in adult and 2.5 in neonates to incorporate the larger ECS (∼15% and ∼40% of the ICS in adults and neonates respectively) observed in neonates [23, 24]. To see how the relative volume of ECS affects the behavior of spontaneous Na^+^ fluctuations, we vary *β* over a wide range in some simulations of neonate network. We remark that using *β* = 2.5 in the network representing the adult brain (mature inhibition) didn’t cause spontaneous Na^+^ fluctuations (not shown). *γ =* 3 × 10^*4*^/(*F* × *r*_*in*_) is the conversion factor from current units to flux units, where *F* and *r*_*in*_ are the Faraday’s constant and radius of the neuron, respectively. The factor 2 in front of *I*_*pump*_ is due to the fact that the Na^+^/K^+^ pump extrudes two K^+^ in exchange for three Na^+^.

The rate of change of [Na^+^]_*i*_ is controlled by *I*_*Na*_ and *I*_*pump*_ [27], that is

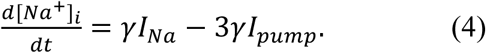

The equations modeling *I*_*pump*_, *I*_*glia*_, and *I*_*diff1*_ are given as

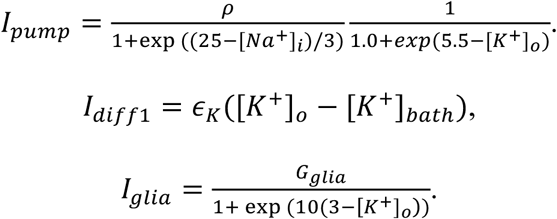

Where Ρ is the pump strength and is a function of available oxygen concentration in the tissue ([O_2_]) or perfusion solution [70], that is

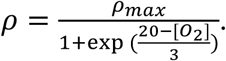

and *ρ*_*max*_, *G*_*glia*_, *ϵ*_*k*_, and [*K*^+^]_*bath*_ represent the maximum Na^+^/K^+^ pump strength, maximum glial *K*^+^ uptake, constant for K^+^ diffusion to vasculature or bath solution, and K^+^ concentration in the perfusion solution respectively. The change in oxygen concentration is given by the following rate equation [70].

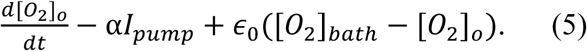

Where [*O*_*2*_]_*bath*_ is the bath oxygen concentration in the perfusion solution, α converts flux through Na^+^/K^+^ pumps (mM/sec) to the rate of oxygen concentration change (mg/(L×sec)), and ε_*O*_ is the diffusion rate constant for oxygen from bath solution to the neuron. We also incorporate lateral diffusion of K^+^ (I_diff2_) between adjacent neurons where the extracellular K^+^ of each neuron in the excitatory layer diffuses to/from the nearest neighbors in the same layer and one nearest neuron in the inhibitory layer. That is,

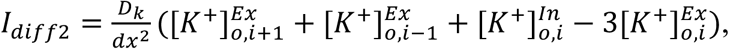

where the subscript *i* indicates the index of the neuron with which the exchange occurs, *D*_*k*_ is the diffusion coefficient of K^+^, and *dx* represents the separation between neighboring cells. The diffusion of K^+^ in the inhibitory layer is modified so that each inhibitory neuron exchanges K^+^ with the two nearest neighbors in the same layer and five nearest neighbors in the excitatory layer. The separation between neighboring neurons in the inhibitory layer is five times that of neighboring neurons in the excitatory layer.

To simplify the formalism, [K^+^]_*i*_ and [Na^+^]_*o*_ are linked to [Na^+^]_*i*_ as previously described [27, 56, 57, 71, 72].

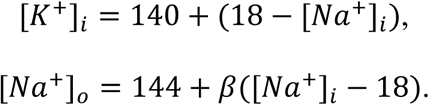

[Cl^−^]_*i*_ and [C^−^l]_*o*_ are given by the conservation of charge inside and outside the cell respectively.

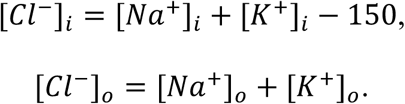

The number 150 in the above equation represents the concentration of impermeable cations. The values of various parameters used in the model are given in Table 1.

**Table 1:** Values and meanings of various parameters used in the model.

| Parameter | Value and Unit (Excitatory, Inhibitory neuron) | Description |
| --- | --- | --- |
| C | 1.0 $\mu\text{F}/\text{cm}^2$ | Membrane capacitance |
| $\gamma$ | $3 \times 10^4/(F \times r_{in})$ mM/(cm$\bullet$ $\mu\text{A}$) | Current to concentration conversion factor |
| $r_{in}$ | 6 $\mu\text{m}$ | Radius of the neuron |
| $\beta$ | 2.5 | Ratio of ICS to ECS |
| $G_{Cl}^L$ | 0.001 mS/cm$^2$ | Conductance of Cl$^-$ leak channels |
| $G_{Na}^F$ | 165 mS/cm$^2$, 35 mS/cm$^2$ | Maximal conductance of fast Na$^+$ channels |
| $G_K^{DR}$ | 80 mS/cm$^2$, 9 mS/cm$^2$ | Maximal conductance of delayed rectified K$^+$ channels |
| $G_K^L$ | 0.02 mS/cm$^2$ | Conductance of K$^+$ leak channels |
| $G_{Na}^L$ | 7.6 $\mu\text{S}/\text{cm}^2$, 8.55 $\mu\text{S}/\text{cm}^2$ | Conductance of Na$^+$ channels |
| $G_{\text{AMPA/NMDA}}$ | 1 $\mu\text{S}/\text{cm}^2$ | Maximal conductance of E-to-E and E-to-I synapses |
| $G_{GABA}^{ii}$ | 10 $\mu\text{S}/\text{cm}^2$ | Maximal conductance of I-to-I synapses |
| $G_{GABA}^{ie}$ | 0.1 - 3.0 mS/cm$^2$ | Maximal conductance of I-to-E synapses |
| $\tau_R$ | 0.1 ms | Rise constant for synaptic gating |
| $\tau_D$ | 4.0 ms, 30.0 ms | Decay constant for synaptic gating |
| $f_{\text{stoch}}$ | 1 Hz, 0.1 Hz | Mean arrival frequency of stochastic input |
| $\rho_{\text{max}}$ | 29 mM/s | Maximum Na$^+$/K$^+$ pump strength |
| [O$_2$]$_{\text{bath}}$ | 32 mg/l | Oxygen concentration in the bath solution |
| $\alpha$ | 5.3 g/mol | Conversion factor from Na$^+$/K$^+$-ATPase current to oxygen consumption rate |
| $\epsilon_o$ | 0.17 s$^{-1}$ | Oxygen diffusion constant |
| $G_{\text{glia}}$ | 60 mM/s | Maximum glia K$^+$ uptake |
| $\epsilon_K$ | 3 s$^{-1}$ | Constant for K$^+$ diffusion between ECS and bath solution (blood vassals *in vivo*) |
| [K$^+$]$_{\text{bath}}$ | 3.0 mM | K$^+$ concentration in the bath solution *in vitro* or in vasculature *in vivo* |
| $D_K$ | $2.5 \times 10^{-5}$ cm$^2$/s | Diffusion coefficient of K$^+$ in the ECS |
| dx | 200 $\mu\text{m}$ | Distance between adjacent neurons |

### Numerical Methods

The rate equations were solved in Fortran 90 using the midpoint method, with a time step of 0.02 ms. The statistical analysis of the data obtained from simulations is performed in Matlab. Codes reproducing key results are available upon request from authors. Significance was determined using students t-tests (p>0.001: ***).

## Acknowledgements

This works was supported by NIH through grant number R01AG053988 (GU), the Deutsche Forschungsgemeinschaft through grant number SPP1757:Ro2327/8-2 (CRR), and a start-up fund of the SPP1757 (LF).

## References

1. Spitzer NC. Electrical activity in early neuronal development. Nature. 2006;444(7120):707–12.

2. Garaschuk O, Linn J, Eilers J, Konnerth A. Large-scale oscillatory calcium waves in the immature cortex. Nature neuroscience. 2000;3(5):452–9.

3. Leinekugel X, Medina I, Khalilov I, Ben-Ari Y, Khazipov R. Ca2+ oscillations mediated by the synergistic excitatory actions of GABAA and NMDA receptors in the neonatal hippocampus. Neuron. 1997;18(2):243–55.

4. Spitzer NC. Spontaneous Ca2+ spikes and waves in embryonic neurons: signaling systems for differentiation. Trends in neurosciences. 1994;17(3):115–8.

5. Felix L, Ziemens D, Seifert G, Rose CR. Spontaneous Ultraslow Na+ Fluctuations in the Neonatal Mouse Brain. Cells. 2020;9(1):102.

6. Katz LC, Shatz CJ. Synaptic activity and the construction of cortical circuits. Science. 1996;274(5290):1133–8.

7. Garaschuk O, Hanse E, Konnerth A. Developmental profile and synaptic origin of early network oscillations in the CA1 region of rat neonatal hippocampus. The Journal of physiology. 1998;507(1):219–36.

8. Ben-Ari Y, Cherubini E, Corradetti R, Gaiarsa J. Giant synaptic potentials in immature rat CA3 hippocampal neurones. The Journal of physiology. 1989;416(1):303–25.

9. Penn AA, Riquelme PA, Feller MB, Shatz CJ. Competition in retinogeniculate patterning driven by spontaneous activity. Science. 1998;279(5359):2108–12.

10. Luhmann HJ, Sinning A, Yang J-W, Reyes-Puerta V, Stüttgen MC, Kirischuk S, et al. Spontaneous neuronal activity in developing neocortical networks: from single cells to large-scale interactions. Frontiers in neural circuits. 2016;10:40.

11. Griguoli M, Cherubini E. Early correlated network activity in the hippocampus: its putative role in shaping neuronal circuits. Frontiers in cellular neuroscience. 2017;11:255.

12. Ben-Ari Y, Khazipov R, Leinekugel X, Caillard O, Gaiarsa J-L. GABAA, NMDA and AMPA receptors: a developmentally regulatedmenage a trois’. Trends in neurosciences. 1997;20(11):523–9.

13. Lohmann C, Kessels HW. The developmental stages of synaptic plasticity. The Journal of physiology. 2014;592(1):13–31.

14. Cherubini E, Gaiarsa JL, Ben-Ari Y. GABA: an excitatory transmitter in early postnatal life. Trends in neurosciences. 1991;14(12):515–9.

15. Ben-Ari Y, Gaiarsa J-L, Tyzio R, Khazipov R. GABA: a pioneer transmitter that excites immature neurons and generates primitive oscillations. Physiological reviews. 2007;87(4):1215–84.

16. Ben-Ari Y, Holmes GL. The multiple facets of γ-aminobutyric acid dysfunction in epilepsy. Current opinion in neurology. 2005;18(2):141–5.

17. Rivera C, Voipio J, Kaila K. Two developmental switches in GABAergic signalling: the K+–Cl− cotransporter KCC2 and carbonic anhydrase CAVII. The Journal of physiology. 2005;562(1):27–36.

18. Kirmse K, Kummer M, Kovalchuk Y, Witte OW, Garaschuk O, Holthoff K. GABA depolarizes immature neurons and inhibits network activity in the neonatal neocortex in vivo. Nature communications. 2015;6(1):1–13.

19. Larsen BR, Stoica A, MacAulay N. Developmental maturation of activity-induced K+ and pH transients and the associated extracellular space dynamics in the rat hippocampus. The Journal of Physiology. 2019;597(2):583–97.

20. MacAulay N. Molecular mechanisms of K+ clearance and extracellular space shrinkage— Glia cells as the stars. Glia. 2020.

21. Felix L, Stephan J, Rose CR. Astrocytes of the early postnatal brain. European Journal of Neuroscience. 2020;in press.

22. Safiulina VF, Zacchi P, Taglialatela M, Yaari Y, Cherubini E. Low expression of Kv7/M channels facilitates intrinsic and network bursting in the developing rat hippocampus. The Journal of physiology. 2008;586(22):5437–53.

23. Lehmenkühler A, Syková E, Svoboda J, Zilles K, Nicholson C. Extracellular space parameters in the rat neocortex and subcortical white matter during postnatal development determined by diffusion analysis. Neuroscience. 1993;55(2):339–51.

24. Nicholson C, Hrabětová S. Brain extracellular space: the final frontier of neuroscience. Biophysical journal. 2017;113(10):2133–42.

25. Zuzana S, Syková E. Diffusion heterogeneity and anisotropy in rat hippocampus. Neuroreport. 1998;9(7):1299–304.

26. McBain CJ, Traynelis SF, Dingledine R. Regional variation of extracellular space in the hippocampus. Science. 1990;249(4969):674–7.

27. Cressman JR, Ullah G, Ziburkus J, Schiff SJ, Barreto E. The influence of sodium and potassium dynamics on excitability, seizures, and the stability of persistent states: I. Single neuron dynamics. Journal of computational neuroscience. 2009;26(2):159–70.

28. Panayiotopoulos C. Neonatal seizures and neonatal syndromes. The Epilepsies: Seizures, Syndromes and Management: Bladon Medical Publishing; 2005.

29. Zanelli S, Rajasekaran K, Grosenbaugh D, Kapur J. Increased excitability and excitatory synaptic transmission during in vitro ischemia in the neonatal mouse hippocampus. Neuroscience. 2015;310:279–89.

30. Van Zundert B, Peuscher MH, Hynynen M, Chen A, Neve RL, Brown RH, et al. Neonatal neuronal circuitry shows hyperexcitable disturbance in a mouse model of the adult-onset neurodegenerative disease amyotrophic lateral sclerosis. Journal of Neuroscience. 2008;28(43):10864–74.

31. Bender RA, Baram TZ. Epileptogenesis in the developing brain: what can we learn from animal models? Epilepsia. 2007;48:2–6.

32. Hauser WA. Seizure disorders: the changes with age. Epilepsia. 1992;33:6–14.

33. Hauser WA. The prevalence and incidence of convulsive disorders in children. Epilepsia. 1994;35:S1–S6.

34. Dzhala VI, Staley KJ. Excitatory actions of endogenously released GABA contribute to initiation of ictal epileptiform activity in the developing hippocampus. Journal of Neuroscience. 2003;23(5):1840–6.

35. Isaev D, Isaeva E, Khazipov R, Holmes GL. Anticonvulsant action of GABA in the high potassium–low magnesium model of ictogenesis in the neonatal rat hippocampus in vivo and in vitro. Journal of neurophysiology. 2005;94(4):2987–92.

36. Khazipov R, Khalilov I, Tyzio R, Morozova E, Ben-Ari Y, Holmes GL. Developmental changes in GABAergic actions and seizure susceptibility in the rat hippocampus. European Journal of Neuroscience. 2004;19(3):590–600.

37. Ben-Ari Y, Holmes GL. Effects of seizures on developmental processes in the immature brain. The Lancet Neurology. 2006;5(12):1055–63.

38. Barger Z, Easton CR, Neuzil KE, Moody WJ. Early network activity propagates bidirectionally between hippocampus and cortex. Developmental neurobiology. 2016;76(6):661–72.

39. Pelkey KA, Chittajallu R, Craig MT, Tricoire L, Wester JC, McBain CJ. Hippocampal GABAergic inhibitory interneurons. Physiological reviews. 2017;97(4):1619–747.

40. Rivera C, Voipio J, Payne JA, Ruusuvuori E, Lahtinen H, Lamsa K, et al. The K+/Cl− co-transporter KCC2 renders GABA hyperpolarizing during neuronal maturation. Nature. 1999;397(6716):251–5.

41. Achilles K, Okabe A, Ikeda M, Shimizu-Okabe C, Yamada J, Fukuda A, et al. Kinetic properties of Cl− uptake mediated by Na+-dependent K+-2Cl− cotransport in immature rat neocortical neurons. Journal of Neuroscience. 2007;27(32):8616–27.

42. Kaila K, Price TJ, Payne JA, Puskarjov M, Voipio J. Cation-chloride cotransporters in neuronal development, plasticity and disease. Nature Reviews Neuroscience. 2014;15(10):637.

43. Bordey A, Sontheimer H. Postnatal development of ionic currents in rat hippocampal astrocytes in situ. Journal of Neurophysiology. 1997;78(1):461–77.

44. Kriegstein A, Alvarez-Buylla A. The glial nature of embryonic and adult neural stem cells. Annual review of neuroscience. 2009;32:149–84.

45. Wang DD, Bordey A. The astrocyte odyssey. Progress in neurobiology. 2008;86(4):342–67.

46. Privat A. Postnatal gliogenesis in the mammalian brain. Int Rev Cytol. 1975;40(28):l-323.

47. Schreiner AE, Durry S, Aida T, Stock MC, Rüther U, Tanaka K, et al. Laminar and subcellular heterogeneity of GLAST and GLT-1 immunoreactivity in the developing postnatal mouse hippocampus. Journal of Comparative Neurology. 2014;522(1):204–24.

48. Syková E, Nicholson C. Diffusion in brain extracellular space. Physiological reviews. 2008;88(4):1277–340.

49. Khalilov I, Le Van Quyen M, Gozlan H, Ben-Ari Y. Epileptogenic actions of GABA and fast oscillations in the developing hippocampus. Neuron. 2005;48(5):787–96.

50. Close B, Banister K, Baumans V, Bernoth EM, Bromage N, Bunyan J, et al. Recommendations for euthanasia of experimental animals: Part 2. DGXT of the European Commission. Laboratory animals. 1997;31(1):1-32. Epub 1997/01/01. doi: 10.1258/002367797780600297. PubMed PMID: 9121105.

51. Gerkau NJ, Lerchundi R, Nelson JS, Lantermann M, Meyer J, Hirrlinger J, et al. Relation between activity-induced intracellular sodium transients and ATP dynamics in mouse hippocampal neurons. The Journal of physiology. 2019;597(23):5687–705.

52. Kafitz KW, Meier SD, Stephan J, Rose CR. Developmental profile and properties of sulforhodamine 101—Labeled glial cells in acute brain slices of rat hippocampus. Journal of neuroscience methods. 2008;169(1):84–92.

53. Langer J, Gerkau NJ, Derouiche A, Kleinhans C, Moshrefi-Ravasdjani B, Fredrich M, et al. Rapid sodium signaling couples glutamate uptake to breakdown of ATP in perivascular astrocyte endfeet. Glia. 2017;65(2):293–308.

54. Langer J, Rose CR. Synaptically induced sodium signals in hippocampal astrocytes in situ. The Journal of physiology. 2009;587(24):5859–77.

55. Kopell N, Börgers C, Pervouchine D, Malerba P, Tort A. Gamma and theta rhythms in biophysical models of hippocampal circuits. Hippocampal microcircuits: Springer; 2010. p. 423–57.

56. Ullah G, Schiff SJ. Assimilating seizure dynamics. PLoS computational biology. 2010;6(5):e1000776.

57. Ullah G, Wei Y, Dahlem MA, Wechselberger M, Schiff SJ. The role of cell volume in the dynamics of seizure, spreading depression, and anoxic depolarization. PLoS computational biology. 2015;11(8):e1004414.

58. Krishnan GP, González OC, Bazhenov M. Origin of slow spontaneous resting-state neuronal fluctuations in brain networks. Proceedings of the National Academy of Sciences. 2018;115(26):6858–63.

59. Somjen G, Kager H, Wadman W. Computer simulations of neuron-glia interactions mediated by ion flux. Journal of computational neuroscience. 2008;25(2):349–65.

60. Huebel N, Ullah G. Anions govern cell volume: a case study of relative astrocytic and neuronal swelling in spreading depolarization. PloS one. 2016;11(3).

61. Hübel N, Hosseini-Zare MS, Žiburkus J, Ullah G. The role of glutamate in neuronal ion homeostasis: A case study of spreading depolarization. PLoS computational biology. 2017;13(10):e1005804.

62. Ermentrout GB, Kopell N. Fine structure of neural spiking and synchronization in the presence of conduction delays. Proceedings of the National Academy of Sciences. 1998;95(3):1259–64.

63. Traub RD, Miles R. Neuronal networks of the hippocampus: Cambridge University Press; 1991.

64. Tort AB, Rotstein HG, Dugladze T, Gloveli T, Kopell NJ. On the formation of gamma-coherent cell assemblies by oriens lacunosum-moleculare interneurons in the hippocampus. Proceedings of the National Academy of Sciences. 2007;104(33):13490–5.

65. Wang X-J, Buzsáki G. Gamma oscillation by synaptic inhibition in a hippocampal interneuronal network model. Journal of neuroscience. 1996;16(20):6402–13.

66. Saraga F, Wu C, Zhang L, Skinner F. Active dendrites and spike propagation in multicompartment models of oriens-lacunosum/moleculare hippocampal interneurons. The Journal of physiology. 2003;552(3):673–89.

67. Krishnan GP, Bazhenov M. Ionic dynamics mediate spontaneous termination of seizures and postictal depression state. Journal of Neuroscience. 2011;31(24):8870–82.

68. Fröhlich F, Bazhenov M, Iragui-Madoz V, Sejnowski TJ. Potassium dynamics in the epileptic cortex: new insights on an old topic. The Neuroscientist. 2008;14(5):422–33.

69. Ullah G, Cressman Jr JR, Barreto E, Schiff SJ. The influence of sodium and potassium dynamics on excitability, seizures, and the stability of persistent states: II. Network and glial dynamics. Journal of computational neuroscience. 2009;26(2):171–83.

70. Wei Y, Ullah G, Ingram J, Schiff SJ. Oxygen and seizure dynamics: II. Computational modeling. Journal of neurophysiology. 2014;112(2):213–23.

71. Wei Y, Ullah G, Schiff SJ. Unification of neuronal spikes, seizures, and spreading depression. Journal of Neuroscience. 2014;34(35):11733–43.

72. Hübel N, Andrew RD, Ullah G. Large extracellular space leads to neuronal susceptibility to ischemic injury in a Na+/K+ pumps–dependent manner. Journal of computational neuroscience. 2016;40(2):177–92.

